# Top-down control of the left visual field bias in cued visual spatial attention

**DOI:** 10.1101/2022.02.02.478855

**Authors:** Sreenivasan Meyyappan, Abhijit Rajan, George R Mangun, Mingzhou Ding

## Abstract

A left visual field (LVF) bias in perceptual judgements, response speed and discrimination accuracy are well documented in humans. However, LVF bias can be modulated by perceptual and task demands. For example, cuing spatial attention can reduce or eliminate the LVF bias, suggesting that attentional control can compensate for the LVF bias. We investigated this possibility directly by recording pupillometry together with fMRI in a cued visual spatial attention task. Prior to the onset of a task-relevant target stimulus, we observed that the pupil was significantly more dilated following attend-right than attend-left cues even though task performance did not differ. This difference in pupil dilation was inversely related to the corresponding difference in later target-evoked pupil dilation and in the reaction times to those targets, suggesting that an increased attentional effort was triggered by the attend-right cues, and this offset the LVF bias, equating behavioral performance. The differences in pupil dilation to the right versus left hemifield were correlated with corresponding fMRI differences primarily in the right hemisphere, supporting the idea that the increased attentional effort for rightward attention is mediated by activity in right hemisphere networks, which illuminates how attentional control mediates attentional biases in vision.

## Introduction

In humans, most models of visual-spatial attention hold that the left and right hemispheres of the human brain differ in their contributions to the control of attention (Kinsbourne 1970; Heilman and Abell 1980; Mesulam 1981). Evidence in support of such models comes from studies in patients with unilateral brain damage, where right hemisphere lesions lead to more severe neglect of contralateral left visual field stimuli than do left hemisphere lesions for right visual field stimuli (Mesulam, 1981; Heilman et al., 1987; Beis et al., 2004; Becker and Karnath, 2007; Duecker and Sack, 2015), and studies in commissurotomy patients, where the two cerebral hemispheres are disconnected at the cortical level and can be investigated separately, revealing differences in left and right hemisphere attention performance (Mangun et al., 1994; Kingstone et al, 1995). In healthy, intact individuals, functional brain imaging and neurophysiological recordings have also shown differences in brain activity between the two hemispheres in attention tasks (Corbetta and Shulman, 2011; Gallotto et al., 2020).

A classic measure of attention biases in neuropsychology is the line bisection task. Patients are asked to bisect horizontal lines on a sheet of paper by drawing a vertical marker in the middle of the line, bisecting it. Patients with right hemisphere lesions tend to bias their line bisections to the right of the horizontal line’s midline, as though they were neglecting the contralateral left portions of the line, and thence misjudging its midline. Interestingly, however, when neurologically intact participants are asked to perform the line bisection task, Bowers and Heilman (1980) observed the opposite tendency: neurologically intact participants tended to place the bisection marker to the left of the midline of the horizontal line, as though the healthy subjects were neglecting the right visual field portions of the line, and thence misjudging the midline as being to the left of the line’s center. This naturally occurring attentional asymmetry, where the left hemi-space is favored over the right by healthy neurologically intact individuals, is referred to as the left visual field bias, or pseudoneglect, and is thought to reflect a right hemisphere dominance in spatial attention (Heilman and Abell, 1980; Reuter-Lorenz et al., 1990; Benwell et al., 2014).

The left visual field bias has been revealed in many different experimental settings. In visual search, when instructed to identify and respond to target objects amongst distracting items in arrays spread across visual space, the participant often begins the exploration of the visual scene on the left side (Bartolomeo et al., 1994; Gigliotta et al., 2017). In the computer version of the line bisection task, when a pre-bisected line is shown, the participant often indicates the left segment to be longer when the bisection marker is actually at the midpoint (Rueckert and Mcfadden, 2004; Dufour et al., 2007; Thomas and Elias, 2011). In rapid-serial-visual-search (RSVP) tasks, when asked to detect targets in a rapid sequence of stimuli presented at single locations in either the left or the right visual field, the participant often performs better for left visual than the right visual field RSVP task (Evert et al., 2003; Holländer et al., 2005; Śmigasiewicz et al., 2010; Verleger et al., 2013; Śmigasiewicz et al., 2014; Śmigasiewicz et al., 2016).

Interestingly, the left visual field bias is diminished or absent when stimulus presentation is preceded by cues that direct attention to the visual hemifield of the upcoming visual stimulus (e.g., Gitelman et al., 1999; Giesbrecht et al., 2003; McCourt et al., 2005; Wilson et al., 2005; Giesbrecht et al., 2006; Corbetta et al., 2008). In a dual-stream RSVP task, Śmigasiewicz et al. (2016) compared responses to left-visual field stimuli versus right-visual field stimuli with or without having instructional cueing preceding the stimuli. In the absence of cueing, there is again evidence of a left visual field bias, consistent with prior studies (Evert et al., 2003; Holländer et al., 2005; Verleger et al., 2009; Śmigasiewicz et al., 2010). With cueing, however, the difference in behavioral performance between attended right-stimuli and attended left-stimuli was diminished. The mechanism underlying the reduction of the left visual field bias in performance with attention precues, remains to be understood. In line with the prior literature, we posit that the left visual field bias results from an asymmetry in the allocation of attention (i.e., in attentional control), with attention to the right visual field requiring more effort than directing it to the left, and we hypothesize that precuing spatial attention rightward engages additional attentional control effort, which in a compensatory fashion, leads to reductions in the left visual field bias in target stimulus processing.

Testing this hypothesis using behavioral measures alone is challenging because of the lack of a behavioral measure during the foreperiod (cue-to-target period) in tasks that permit the isolation of attentional control from attentional target selection; this is simply because there is no behavioral response until the target appears. Pupillometry offers a potential solution. Past work has shown that pupil diameter is a reliable physiological index of effort (Kahneman and Beatty, 1966; Beatty, 1982; Ebitz and Moore, 2019) and can be used to measure attentional states (Hoeks and Levelt, 1993; Kang et al., 2014; Binda and Murray 2015; Mathôt et al., 2013; Liao et al., 2016; Irons et al., 2017; Wainstein et al., 2017). For example, Irons and colleagues (2017) showed that attentional cues (both auditory and visual) elicit pupil dilation, and this increase in pupil size reflects task difficulty, (i.e., more difficult task conditions elicit larger pupil dilations). If directing and maintaining covert visual spatial attention to the right visual field requires more attentional effort, one would predict that the pupil would be more dilated following attend-right cues than attend-left cues in a spatial cuing paradigm, which in turn would diminish the left visual field bias.

We recorded behavioral and pupillometry data from participants performing a cued visual spatial attention task. To avoid the confounding influences from the pupil’s reflexive responses to visual stimulation, we used auditory cues instead of visual cues to direct attention, thereby ensuring that cue-related pupillary responses were entirely attributable to internally generated attentional processes and not related to changes in visual stimulation. In addition, fMRI data were collected along with behavioral and pupillometry data, with the goal to use the multimodal data to examine the neural basis of the hypothesized differential pupil responses to left versus right attention cues.

## Materials and Methods

### Overview

The experimental protocol was approved by the Institutional Review Board (IRB) of the University of Florida. Twenty (5 female; mean age 24.65 ± 2.87) right-handed healthy individuals provided written informed consent and took part in the study. The participants reported no prior history of mental disorders and had normal or correct-to-normal vision. The data from this experiment has been used in previous publications to investigate different questions (Rajan et al., 2019; Rajan et al., 2021; Meyyappan et al., 2021).

### Procedure

Two sets of dots, 3.6 degrees lateral to the upper left and upper right of the fixation cross, indicated the two locations where the stimuli would appear. See Figure 1 for illustration. At the beginning of each trial, an auditory cue instructed the participants to covertly direct their attention to either a spatial location (“left” or “right”) or a color (“red” or “green”) while fixating the central plus sign. On 80% of such trials, following a random delay period ranging from 3000 ms to 6600 ms, two colored rectangles (red or green) were presented for a duration of 200 ms, with one in each of the two peripheral locations. For the remaining 20% of the trials, the cues were not followed by the stimuli (cue-only trials), and these cue-only trials were included to help with better modeling of and differentiation between cue-evoked vs target-evoked BOLD activities. Following the presentation of the stimuli, the subject’s task was to report the orientation of the rectangle (target) appearing at the cued location or having the cued color, and to ignore the other rectangle (distractor). For color trials, the two rectangles displayed were always of the opposite color; for spatial trials, the two rectangles were either of the same color or of the opposite color. On 8% of the trials (invalid trials), only one rectangle was displayed, which was either not in the cued location for spatial trials or not having the cued color for color trials, and the participants were required to report the orientation of that rectangle. These invalidly cued trials were included to measure the behavioral benefits of attentional cuing (Posner 1980). An inter-trial interval, which was varied randomly from 8000 ms to 12800 ms following the target onset, elapsed before the start of the next trial. In addition to spatial and color cues (80% of all the trials), there was a third type of cue (20% of all the trials) consisting of the word “none”, which informed the subject to prepare to respond to the orientation of the rectangle being placed on a grey patch (neutral attention). Trials were organized into blocks, with each block consisting of 25 trials and lasting approximately seven minutes. Each participant completed 10 to 14 blocks over two days. For this study, given the stated purpose of investigating the possible existence of a left visual field bias in the control of visual spatial attention, only spatial trials were considered.

**Figure 1.**
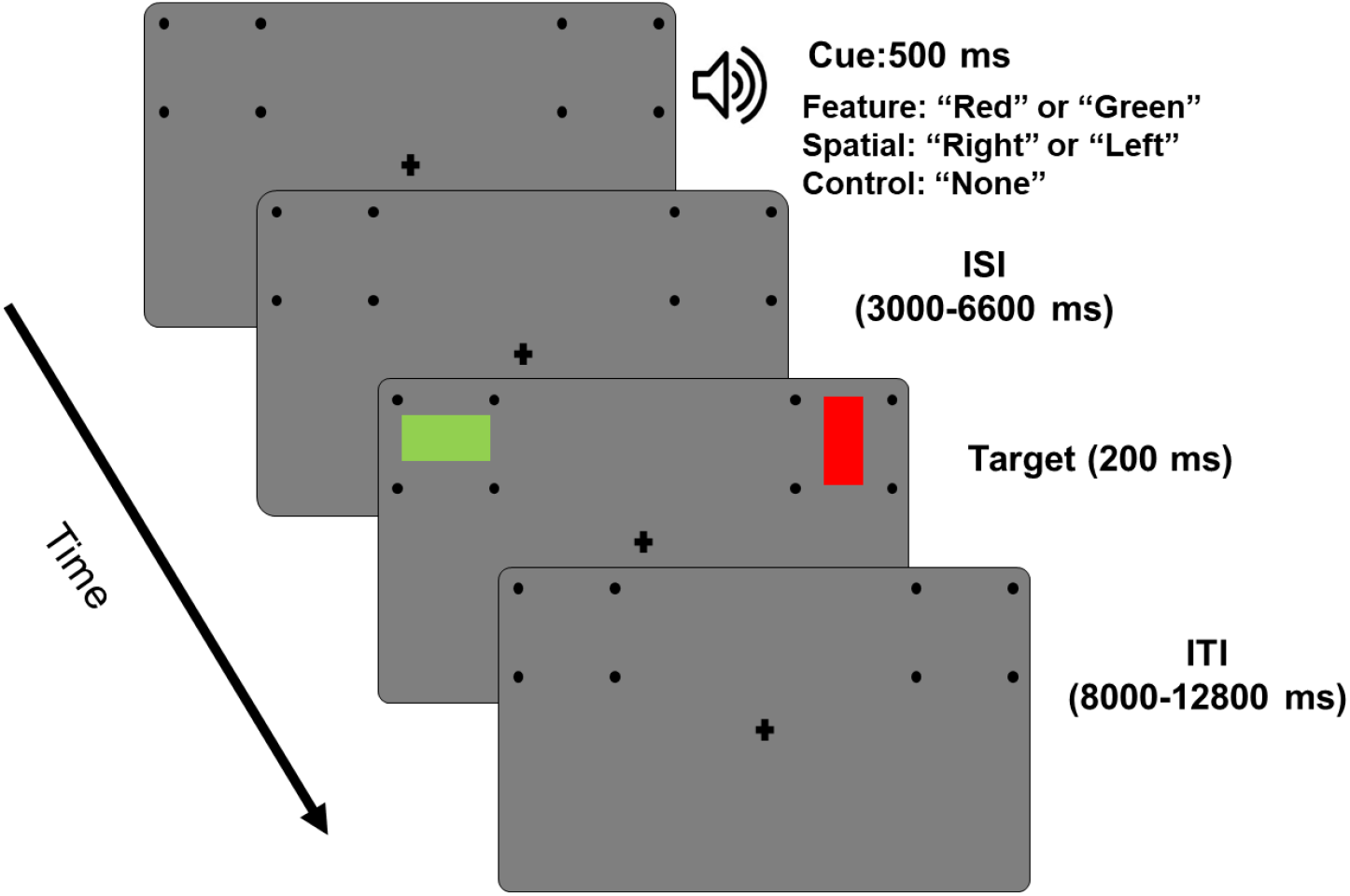
Experimental paradigm for cued feature and spatial attention study. Each trial starts with an auditory cue (500 ms) instructing the subject to covertly attend to a spatial location (“left” or “right”) or to a color (“red” or “green”). Following a variable cue-to-target delay (3000-6600 ms), two colored rectangles were displayed (100 ms), one in each of the two-peripheral location. Participants were asked to report the orientation of the rectangle (horizontal or vertical) displayed in the cued location or having the cued color. On some of the trials (8%) the cues were not valid, i.e., only one target appeared, which was either not at attended location or not having the attended feature, and participants were required to report the orientation of the rectangle. An inter-trial interval, varied randomly from 8000-12800 ms following the target onset, elapsed before the start of the next trial. Only spatial trials were considered in this study.

The participants went through a training session prior to scanning in which they were introduced to the task and became comfortable performing it. Since the study required participants to maintain central fixation for long durations and pay covert attention to the periphery while fixating the central plus sign, they were screened based on their ability to maintain eye fixation throughout the training session. An SR Research EyeLink 1000 eye tracker system was used for that purpose. In addition, participants who attained behavioral accuracy above 70% at the end of the training session proceeded to fMRI-pupillometry recordings.

### Functional MRI Data Acquisition and Preprocessing

Functional magnetic resonance imaging (fMRI) data were collected on a 3T Philips Achieva scanner with a 32-channel head coil (Philips Medical Systems, the Netherlands). The echo-planar imaging (EPI) sequence parameters were: repetition time (TR), 1.98 s; echo time, 30 ms; flip angle, 80°; field of view, 224 mm; slice number, 36; voxel size, 3.5 × 3.5 × 3.5 mm; matrix size, 64 × 64. The slices were oriented parallel to the plane connecting the anterior and posterior commissures.

The fMRI BOLD (Blood Oxygen Level Dependent) data were preprocessed using the Statistical Parametric Mapping toolbox (SPM-12) as well as custom scripts written in MATLAB. Preprocessing steps included slice timing correction, realignment, spatial normalization, and smoothing. Slice timing correction was carried out using sinc interpolation to correct for differences in slice acquisition time within an EPI volume. The images were then spatially realigned to the first image of each session by a 6-parameter rigid body spatial transformation to account for head movement during acquisition. Each participant’s images were then normalized and registered to the Montreal Neurological Institute (MNI) space. All images were further resampled to a voxel size of 3 × 3 × 3 mm, and spatially smoothed using a Gaussian kernel with 7 mm full width at half maximum. Slow temporal drifts in the baseline were removed by applying a high-pass filter with cutoff frequency set at 1/128 Hz.

### Estimating Single Trial BOLD Data

A beta series regression method (Rissman et al., 2004) was used to estimate BOLD response on each trial in every voxel. In this method, cues and targets in trials with correct responses were assigned individual regressors and one regressor was assigned for all the trials with incorrect responses. The regressors were modelled in the conventional GLM framework using custom MATLAB scripts developed within SPM toolbox.

### Pupillometry Data Acquisition and Preprocessing

Pupil diameter data were collected simultaneously with fMRI data using a MR-compatible eye tracker at a sampling rate of 1000Hz (EyeLink 1000, SR Research). During preprocessing, eye blinks were detected, and the pupil diameter during eye blinks was determined by a linear interpolation algorithm (Bradley et al., 2008; Siegle et al., 2008). For cue-related analysis, the continuous pupil data were epoched from 200 ms before cue onset to 3000 ms after cue onset (−200 to 3000ms). The pre-cue (-200 to 0 ms) period was used as a baseline to compute the percentage change of cue-related pupil dilation. Similarly, for target-related analysis, the continuous pupil data were epoched from 200 ms before the target onset to 3000 ms after the target onset. The pre-target (−200 to 0 ms) period was used as a baseline to compute the percentage change of target-related pupil dilation.

### Pupillary Response to Cue and Target

To detect whether pupil diameter time courses following attend-left cues and attend-right cues differed, we applied a non-parametric permutation method, which included the following steps. (1) For every subject, attend-left and attend-right labels were randomly shuffled to yield surrogate datasets. (2) The pupil diameter within the surrogate attend-left and attend-right conditions were averaged, and a mean surrogate-left and-right pupil diameter were computed. (3) A paired t-test was performed between mean surrogate-right and -left pupil diameters. (4) The resulting p values as a function of time were analyzed, the contiguous time periods in which the difference was significantly different at p < 0.05 was determined, and the duration, denoted x, of each of these time periods was recorded. (5) Steps (1) through (4) were repeated 10^5^ times, the distribution of x was created, and a threshold corresponding to p < 0.0001 was ascertained from the null hypothesis distribution. (6) For actual data, the time period in which the pupil diameter difference between attend-left and attend-right trials was significantly different at p < 0.05 was determined, and the duration of the time period compared to the threshold determined in Step (5). (7) Pupil diameter between attend-left and attend-right were considered significantly different if the duration of the time period in which pupil diameter difference was significantly different at p < 0.05 was greater than the threshold duration. A similar protocol was followed for analyzing target-evoked pupil response.

### Pupil versus Behavior

To relate cue-evoked pupil diameter with behavior, the trial-by-trial pupil dilation time-courses were averaged to yield an average pupil dilation time course within each subject, and at the population level, a time period of interest was identified using the statistical analysis detailed above (i.e., 1050-3000 ms post-cue). For inter-subject analysis, the pupil diameter difference between attend right and attend left was computed during this time period of interest for each subject and correlated with behavior across subjects. For intra-subject analysis, recall that the experiment consisted of multiple blocks (~12 blocks per participant), also known as runs. The natural occurrence of behavioral fluctuations across these blocks provided the opportunity to relate pupil dynamics and behavioral performance in an intra-subject manner. Specifically, the reaction time difference between attend right and attend left trials and the corresponding cue evoked pupil diameter difference (1050-3000 ms) were computed for each run. The runs were then sorted based on the RT difference and split (median) into two groups: low RT difference group and high RT difference group. The pupil differences in each of the two groups were computed within each subject and averaged across subjects. A paired t-test was then used to compare the pupil dilation differences estimates between low and high RT difference groups.

### Pupil-BOLD Coupling

The trial-by-trial fMRI beta-series data for attend-right and attend-left trials were subjected to a two sampled t-test to yield a voxel-wise t-statistic for all the subjects. This voxel-wise estimate of fMRI activation difference was then correlated with pupil dilation difference between attend-right and attend-left, across subjects, using the Pearson correlation technique. The p values from the correlation analysis were subjected to cluster thresholding; clusters containing more than 200 continuous voxels of a given p value were then extracted to yield the brain maps.

## Results

### Behavioral Analysis

Reaction time (RT) for attend-left and attend-right trials were 998 ± 150 ms and 1037 ± 169 ms, respectively, with RT for attend-right trials marginally longer than that for attend-left trials (p < 0.09); see Figure 2A. The response accuracy (percentage of correct trials) was comparable between the two attention conditions (attend-left: 94.0 ± 1.2, attend-right: 93.9 ± 1.2; p < 0.95; see Figure 2B). These results are consistent with previous reports that in cued visual spatial attention paradigms, there is no obvious left visual field bias in behavioral performance (Giesbrecht et al., 2003; Liu et al., 2016).

**Figure 2.**
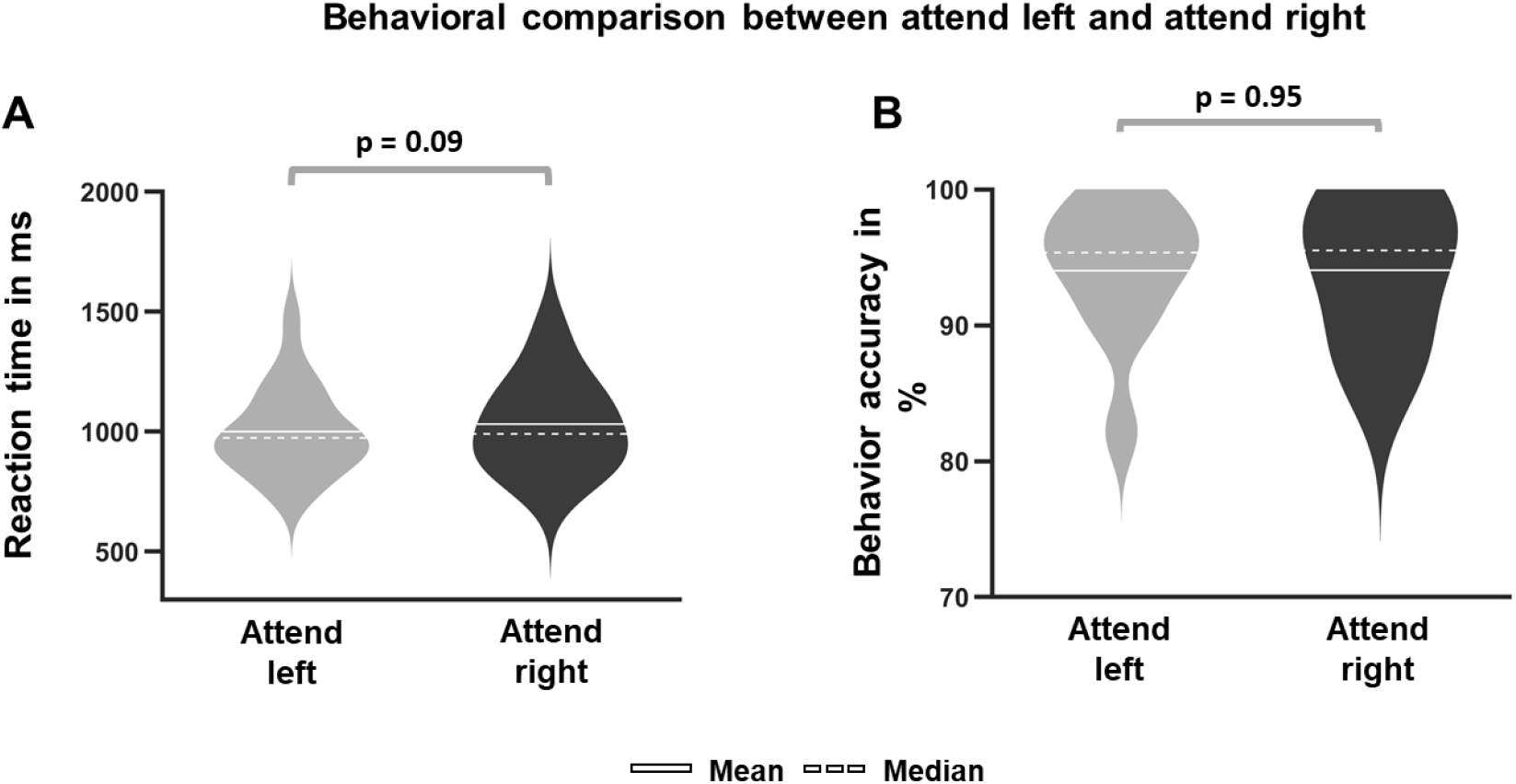
Comparison of behavioral results between spatial attention conditions (attend-right versus attend-left). A) Reaction time. RT for attend-right trials were marginally longer than attend-left trials (p-= 0.09). B) Accuracy. The accuracies were not different between the two types of trials (p = 0.95).

### Cue-Evoked Pupillary Response

As shown in Figure 3A, starting from 1050 ms until the end of the time interval investigated (i.e., 3000 ms post cue), pupil dilation evoked by attend-right cues was significantly larger than that evoked by attend-left cues (attend-right > attend-left; cluster time window threshold p < 0.05), suggesting that attending the right visual field is more effortful than attending the left visual field. To further quantify the effect, as shown in Figure 3B, the pupil dilation for attend-right trials averaged over the interval 1050 to 3000-ms were found to be significantly larger than that for attend-left trials (attend-right > attend-left, p < 0.002). These results support the hypothesis that in covert visual spatial attention there exists a left visual field bias; specifically, the left visual field is favored over the right visual field when deploying attention in advance of stimulus processing and more effort is required to attend to the right visual field to overcome this tendency.

**Figure 3.**
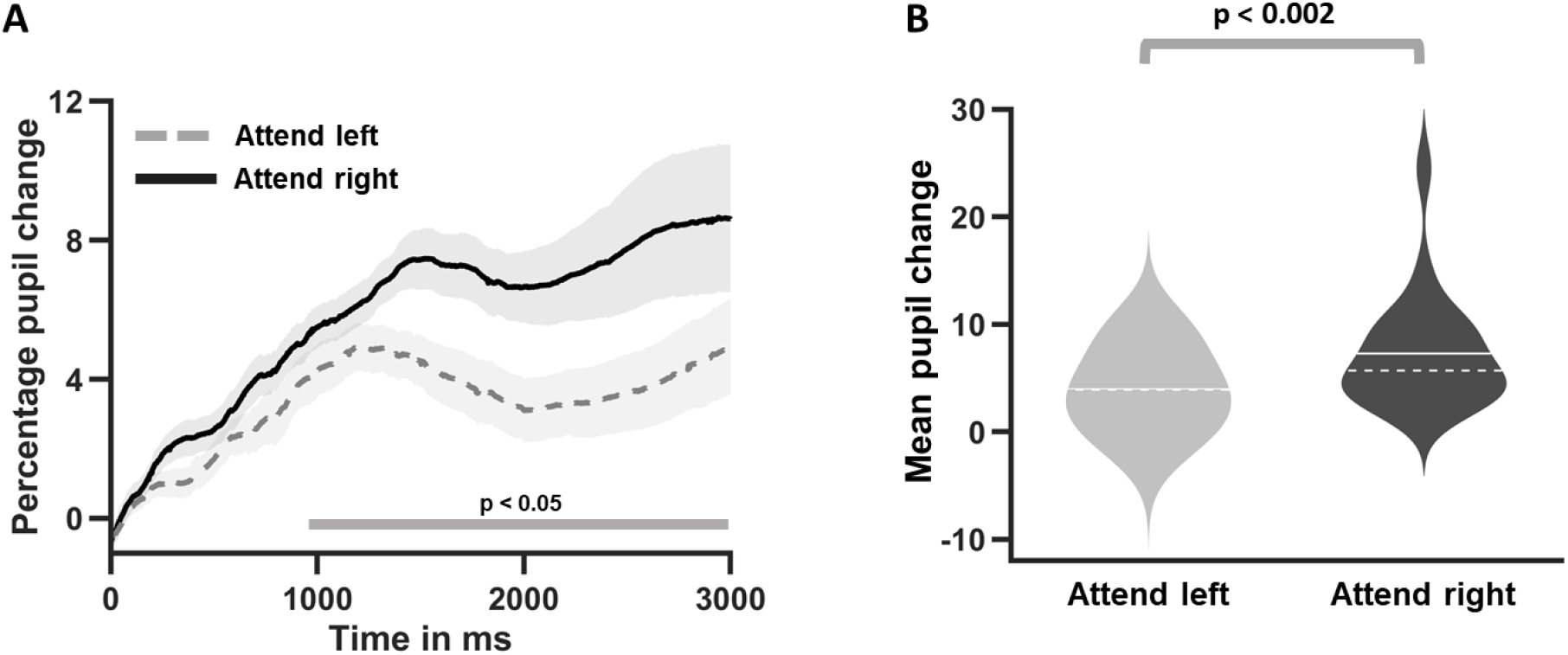
Cue evoked pupillary response during spatial attention trials. A) Time course of pupil dilation measured by the percentage change relative to pre-cue baseline for attend-left and attend-right trials. The shaded region indicates the time window when the difference between attend-left and attend-right trials were significantly different. B) Comparison of cue-evoked pupil dilation averaged over 1050 to 3000 ms interval.

### Cue-Evoked Pupil Response vs Behavior

The increased attentional effort associated with initiating and maintaining anticipatory attention to the right visual field was hypothesized to underlie the disappearance of the left visual field bias in stimulus processing. To test this, we examined the relation between pupil dilation and RT using an intra-subject analysis, which leverages the natural fluctuations of behavioral performance across different experimental runs. Specifically, for each run, we computed the RT difference between attend-right and attend-left trials. Based on this difference, the runs were divided into two groups: high RT difference group and low RT difference group. We then compared the corresponding pupil dilation differences between the two groups.

As shown in Figure 4A and 4B, in the low RT difference group (Figure 4A), the pupil dilation difference between attend-right trials and attend-left trials was significantly greater than 0 in the time period 1300 - 3000 ms post cue onset (attend-right > attend-left, p < 0.05, demonstrating that when there was greater attentional effort associated with attend-right trials (pupil difference being high), the left visual field bias in stimulus processing was diminished (RT difference being low). In contrast, in the high RT difference group (Figure 4B), the pupil difference between attend-right and attend-left trials was not significantly greater than 0 for the entire duration of 0 to 3000 ms, demonstrating that when attentional effort during the cue-target interval was approximately equal between attend-left vs attend-right trials (pupil difference being low), there was a left visual field bias in stimulus processing (RT difference being high). These results support the hypothesis that the increased attentional effort associated with initiating and maintaining covert attention to the right visual field offsets the left visual field bias and equalize performance.

**Figure 4.**
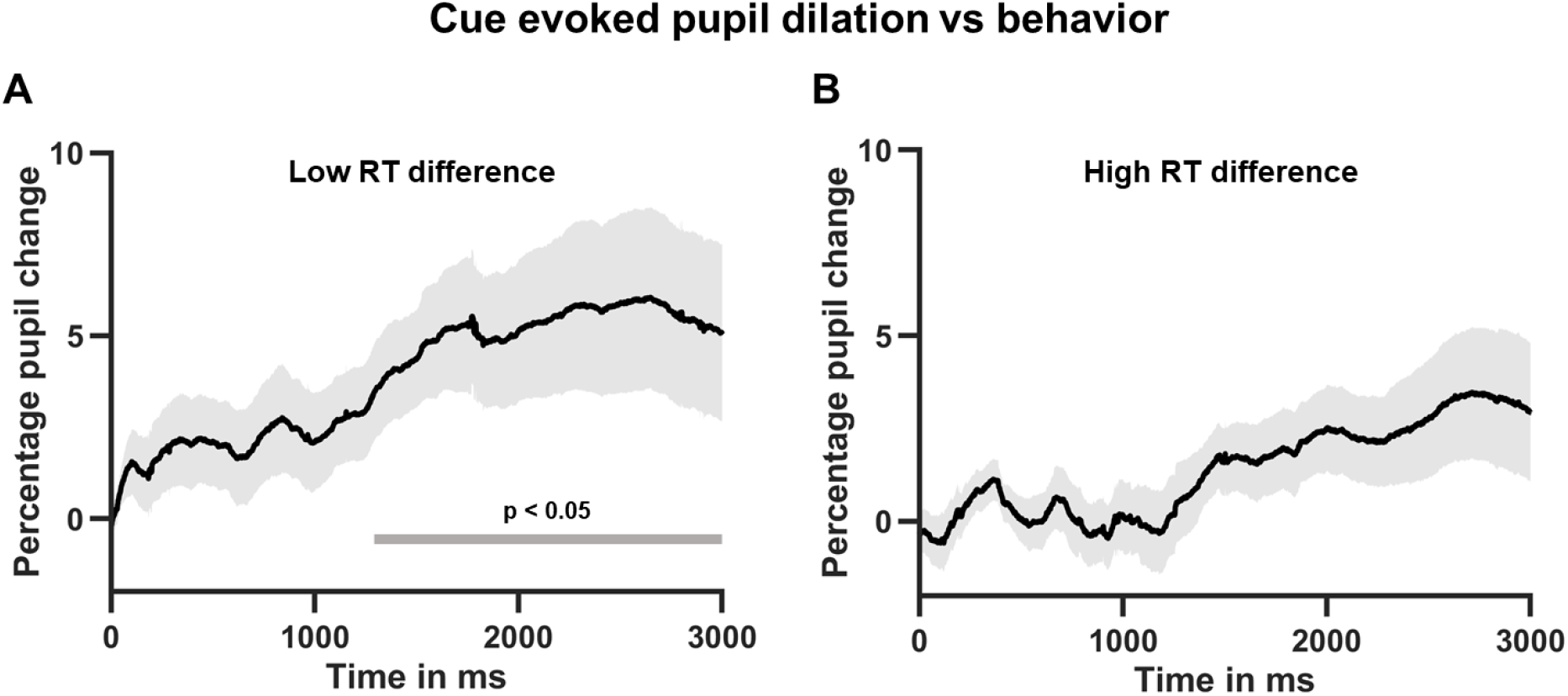
Relationship between cue-evoked pupil dilation and behavior. Experimental runs were sorted into two separate groups: low RT difference group and high RT difference group. A) Timecourse of pupil dilation difference between attend-right and attend-left trials for the low RT difference group. The green horizontal line indicates statistically significant time period. B) Timecourse of pupil dilation difference between attend-right and attend-left trials for the high RT difference group. The curve was not significantly different from zero.

### Relation between Cue-Evoked and Target-Evoked Pupil Responses

The impact of cue-evoked pupil dilation difference on target processing was further examined by relating it to target-evoked pupil dilation difference. As shown in Figure 5A, at the population level, pupil dilation evoked by stimuli during attend-left versus attend-right conditions were not significantly different. Sorting the experimental runs according to the mean target evoked pupil dilation difference (target-right – target-left in the period of 0 to 2000 ms post target) into two separate groups: low target-evoked dilation difference group and high target-evoked dilation difference group, the time-courses of cue-evoked pupil dilation difference (attend-right – attend-left) during attend-right and attend-left trials was computed for the two groups, and the results are shown in Figure 5B and 5C. For the low target-evoked pupil difference group, cue-evoked pupil dilation difference was significantly greater than 0 (attend-right > attend-left; cluster time window threshold p < 0.05) in 1150-3000 ms (Figure 5B), whereas for the high target-evoked pupil difference group, cue-evoked pupil dilation difference was not significantly different from 0 (Figure 5C). The target-evoked pupil analysis, consistent with the RT analysis, again demonstrates that the increased attentional effort in maintaining covert attention to the right visual field helps to diminish the difference in processing targets appearing in left vs right visual field.

**Figure 5.**
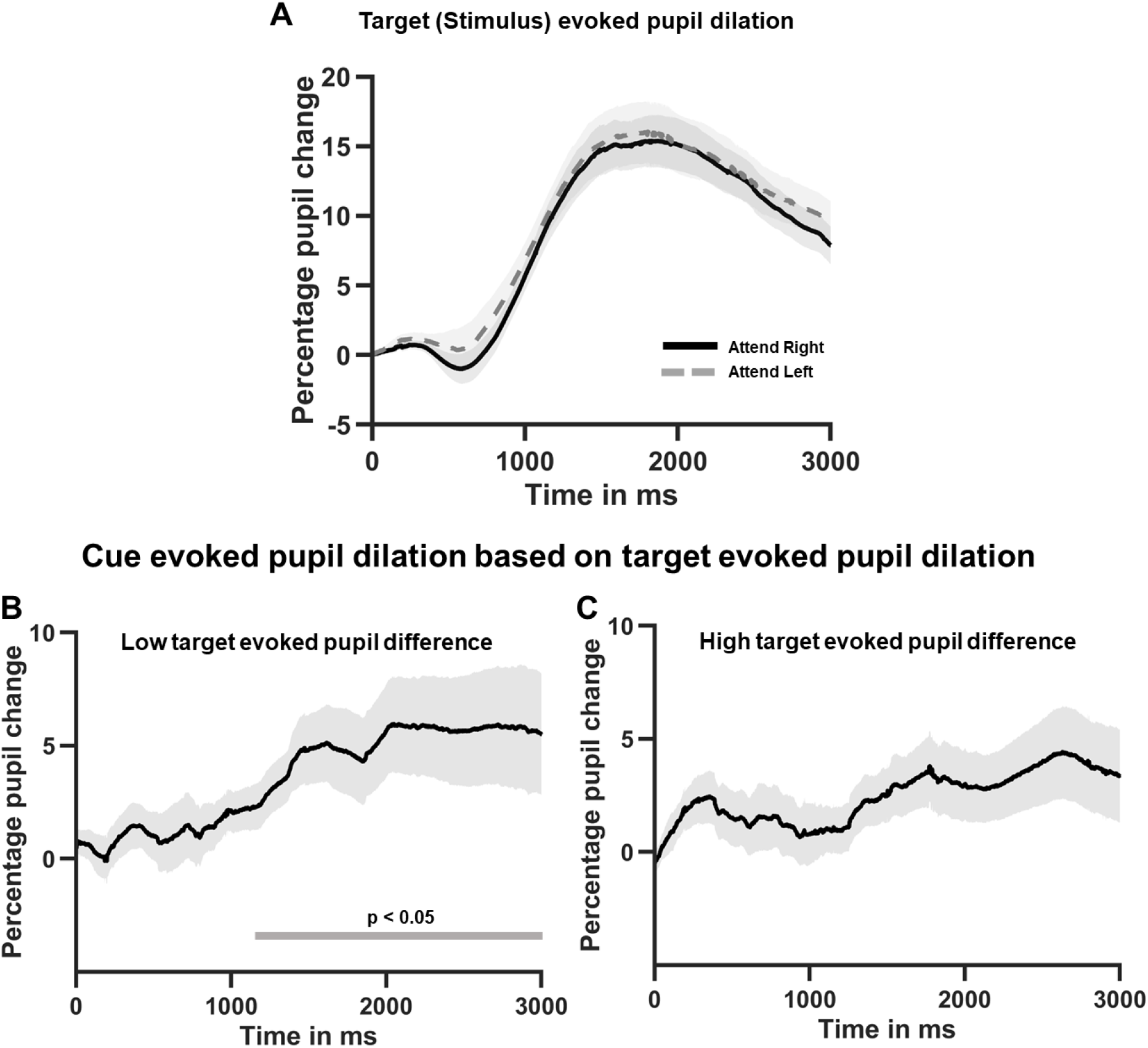
Relationship between target- and cue-evoked pupil size. A) Pupil size over time following bilateral target onset during attend right and left trials. To investigate the relationship between target- and cue-evoked pupil dilation, the subject-level target-evoked pupil differences were sorted into two separate groups: low target-evoked dilation difference group (i.e., no pseudoneglect) and high target-evoked dilation difference group. Panels B and C display the timecourses of cue-evoked pupil dilation difference between attend-left and attend-right conditions for the two target-evoked pupil size groups. B) Timecourse of cue-evoked pupil dilation difference for the low target-evoked dilation difference group; the green line indicates significance. C) Timecourse of cue-evoked pupil dilation difference for the high target-evoked dilation difference group; the timecourse was not significantly different from zero.

### Neural Substrate of Differential Cue-Evoked Pupillary Response

The left visual field bias in stimulus processing is thought to result from the dominant role played by the right hemisphere in spatial cognition (Bowers and Heilman 1980; Benwell et al., 2014). We expected a similar neural substrate for the left visual field bias in stimulus anticipation (cue-to-target interval). To test the hypothesis the pupil difference between attend-left and attend-right in the 1050-3000 ms interval post cue and the corresponding BOLD activity difference were computed at the individual subject level, and then correlated across subjects for all voxels. Voxels showing significant positive correlation at p < 0.05 and being part of a cluster containing 200 or more contiguous such voxels are shown in Figure 6. As expected, more regions in the right hemisphere were positively correlated with pupil dilation difference, including right intra-parietal sulcus (IPS) and right ventro-lateral pre-frontal cortex (VLPFC). See Table 1 for a complete list of areas.

**Table 1.** Regions where differential activities (attend right – attend left) are positively coupled with cue-evoked pupil dilation difference. p < 0.05 (cluster corrected; k = 200).

| Left hemisphere | Right Hemisphere |
| --- | --- |
| Medial superior frontal gyrus | Medial superior frontal gyrus |
| Supplementary motor cortex (SMA) | Supplementary motor cortex (SMA) |
| Paracentral lobule | Intra parietal sulcus (IPS) |
|  | Primary visual cortex |
|  | Occipital Mid |
|  | Inferior temporal cortex |
|  | Ventro-lateral prefrontal cortex (VLPFC) |
|  | Primary motor cortex |
|  | Postcentral gyrus |

**Figure 6.**
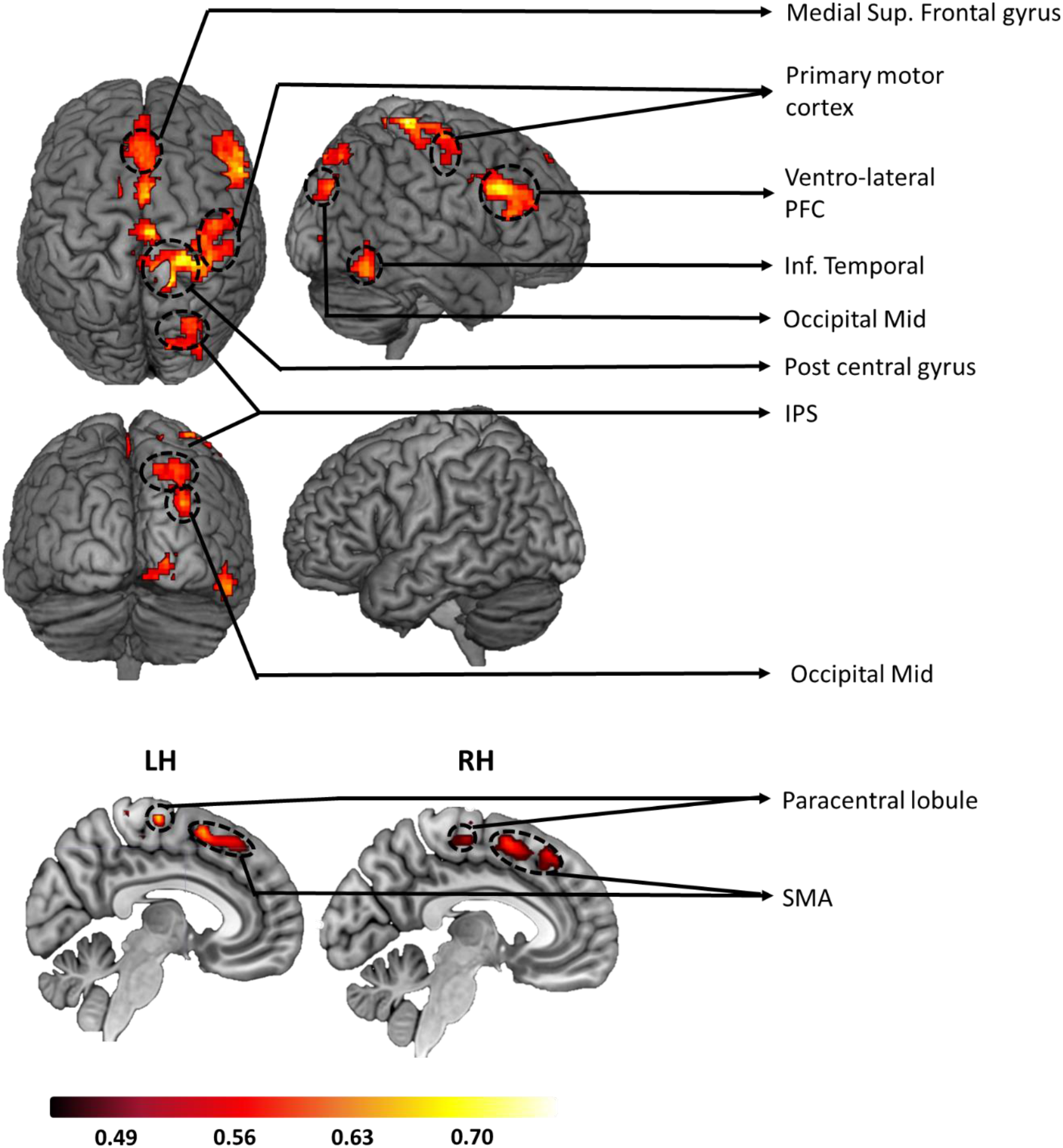
Pupil-BOLD coupling. Regions showing positive coupling between cue-evoked pupil dilation difference and cue-evoked fMRI activation difference between attend-right and attend-left trials. These across-subject correlation maps were thresholded at p < 0.05 (clusters above 200 voxels). Warmer colors reflect stronger correlation coefficients.

## Discussion

We investigated the relationship between attentional control and target processing for leftward and rightward covert visual spatial attention in the context of the well-known left visual field bias in visual processing among healthy human volunteers. Pupillometry was combined with fMRI to provided measures of both attentional effort and the underlying brain networks involved for leftward versus rightward attentional orienting during anticipatory attention. In line with the literature, we posited that there is a left visual field bias in attentional control, with attention to the right visual field requiring more control effort than directing it to the left. We hypothesized that precuing spatial attention rightward engages additional attentional control effort compared to cuing attention leftward; the result of this increased attentional control effort for rightward attention would be to reduce the left visual field bias (pseudoneglect) in subsequent target processing.

We found that attend-right cues evoked a larger dilation of the pupil than attend-left cues, supporting the premise that there is an inherent asymmetry in the effort required for the control of visual spatial attention leftward versus rightward in visual space in response to an attention-directing cue. In addition, trials with larger pupil dilation differences in the cue-target interval were associated with smaller RT and pupil differences time-locked to the appearance of the target. Finally, combining fMRI and pupillometry data, the difference in attentional control effort in the cue-to-target interval between the two types of attention trials was found to be mediated mainly by structures in the right hemisphere, including the right IPS and the right VLPFC.

### Left Visual Field Bias in Stimulus Processing and Attention control

The left visual field bias in stimulus processing (pseudoneglect) has been demonstrated amply since the early 1980s, including in such paradigms as the classic line bisection task and related landmark tasks, visual search tasks, tasks involving processing of lateralized stimuli, and rapid visual serial presentation (RSVP) tasks. This bias is present under circumstances where the physical features and behavioral relevance of stimulus/target events in the left and right visual fields are equivalent. The left field bias can, however, be reduced or reversed when attention is drawn to one visual field via variation in the stimuli used (e.g., McCourt et al., 2005), or the introduction of attention directing cues (Reuter-Lorenz et al., 1990; Hopfinger et al., 2000; Giesbrecht et al., 2003; Śmigasiewicz et al., 2014; Liu et al., 2016); we replicated this general finding here, where neither accuracy nor RTs were significantly different between attend-left and attend-right conditions.

What underlies the cue-related reduction/elimination of the left visual field bias in stimulus processing? We propose, in line with the literature, that there is an inherent asymmetry in attentional control such that in the default state, visual spatial attention favors (is biased toward) the left visual field, but that the top-down attention control system can compensate for this asymmetry. This compensation required additional attentional effort, which is revealed in pupil size. This view is in contrast to prior explanations that emphasized the informativeness of cues in reducing the spatial uncertainty of target location (e.g., Śmigasiewicz et al., 2016). Our ideas were tested using pupillometry (Kahneman and Beatty 1966, Peavler 1974; Kang et al., 2014; Querino et al., 2015). The use of auditory cues in our experimental design allowed us to attribute changes in pupil diameter to attention processes not confounded by reflexive pupil responses to visual cues. The pupil diameter was found to increase following both types of cues, in agreement with previous studies, but importantly, attend-right cues evoked significantly larger pupil dilation compared to attend-left cues, thereby providing support for the idea that attending the right visual field is more effortful than attending the left visual field.

Does the difference in attentional effort in the cue-target interval explain diminished left visual field bias in target processing? We leveraged naturally occurring variability in behavioral performance to examine this question, dividing the runs among the subject population into two groups based upon their RT difference for leftward versus rightward cued attention to the target stimuli. We then looked at the cue-related (pre-target) different in pupil size for each group. The runs having smaller target-related differences in RT, showed larger differences in pupil size in the cue period (attentional control period), and vice versa. Thus, when more attentional effort is expended in response to the attention-directing cues to compensate for the existing left visual bias in attention, there is reduced asymmetry in processing the target stimuli.

In a similar vein, further support for the compensatory attentional control concept comes from the stimulus-evoked pupil dilation data, where we found that smaller stimulus-evoked pupil dilation difference was correlated with larger cue-evoked pupil dilation difference, and vice versa. We interpret these findings to indicate that the increased attentional effort engaged when cues direct attention rightward (observed in cue processing) compensates for the inherent left visual field bias to benefit subsequent target stimulus processing, manifesting as reduced or absent left visual field bias in measures of target processing (RT and pupil size).

### Neural substrate of increased attention effort

Visual spatial attention is controlled by two major brain networks: the dorsal attention network (DAN), comprising bilateral FEF and IPS, is responsible for top-down attention control, whereas the ventral attention network (VAN), comprising right TPJ and right VLPFC, is involved in attentional reorienting (Corbetta and Shulman 2002). Extensive evidence implicates the VAN and right DAN in mediating the left visual field bias in stimulus processing (Fink et al., 2001; Foxe et al., 2003; Çiçek et al., 2009; Benwell et al., 2014). We therefore expected that the increased attentional effort associated with covertly attending the right visual field is also mediated by right hemisphere structures including VAN and right DAN. Correlating pupil dilation difference between attend-right and attend-left attention conditions with the corresponding difference in fMRI activation yielded a set of brain regions mainly in the right hemisphere, including right IPS and right VLPFC, consistent with the prediction. Other regions in the pupil-fMRI correlation map include the lingual gyrus and occipital cortex. These regions have been reported in other pupil-fMRI correlation studies (DiNuzzo et al., 2019).

### Summary

In this study, a left visual field bias in anticipatory visual spatial attention was reported, where pupil was more dilated when subjects were cued to covertly attend right than left visual space in order to prepare to discriminate an upcoming lateralized target stimulus. This asymmetry in the allocation of spatial attention control effort between the visual hemifields was inversely correlated with attentional biases (pseudoneglect) in behavioral measures of stimulus processing and were correlated with right hemisphere brain activity. We interpret these findings as supporting a compensatory control model of visuospatial attention asymmetries that is mediated by right hemisphere networks involved in attentional control.

## Acknowledgements

This work was supported by NIMH grant MH117991 to George R. Mangun and Mingzhou Ding. All data will be publicly available on the NIMH Data Archive.

## Notes

### Competing Interest Statement

The authors have declared no competing interest.

